# Evaluation method of ecosystem service value under complex ecological environment: A case study of Gansu Province, China

**DOI:** 10.1101/2020.09.24.311217

**Authors:** Xiaojiong Zhao, Jian Wang, Junde Su, Wei Sun

## Abstract

Scientific assessment of regional ecosystem service value (ESV) is helpful to make scientific ecological protection plan and compensation policy. The evaluation method has not been established that is adapted to the complex and diverse characteristics of the ecological environment. This paper takes Gansu Province as an example, on the basis of fully considering the regional differences of ecosystem service function, the five correction index of value equivalent factor per unit area were constructed in the provincial scale, and the regional difference adjustment index of 11 kinds of ES was constructed in regional scale, in the way, the value evaluation model based on regional difference was established. The results show that in 2015, the total ESV reached 2239.555 billion Yuan in Gansu Province, ESV gradually increased from the northeast to the southwest, and the high-value areas of service function located in Qilian Mountain and Longnan Mountain, of which the forest and grassland ecosystem contributed the most to the value. From the perspective of value composition, local climate regulation and biodiversity maintenance function are the main service functions of Gansu Province. From 2000 to 2015, ESV increased by 3.426 billion Yuan in Gansu Province, the value of forest and urban ecosystem continued increasing, while the value of cultivated land ecosystem continue decreasing. From the spatial characteristics of service value change, the area of value reduction gradually moved from the central part of Gansu Province to the surrounding area. In general, although this study needs further improvement, the constructed evaluation method provides a relatively comprehensive evaluation scheme for the spatiotemporal dynamic evaluation of ESV in Gansu Province. This study provided a more overall research idea for the evaluation of ESV under complex ecological environment.

## Introduction

Ecosystems can not only provide various raw materials or products directly for human survival, but also have functions such as regulating climate, purifying pollution, conserving water sources, maintaining water and soil, preventing wind and sand, reducing disasters, and protecting biodiversity. All ecosystem products and services are collectively referred to as ecosystem services (ES) [1,2].The evaluation of ecosystem service value (ESV) is the basis of regional ecological construction, ecological protection, ecological work division and ecological decision-making of natural assets, which has become a research hotspot of Ecology[3-5].Since Costanza first accounted for the value of global ES in 1997, the ESV calculation has been gradually used as the core basis of ecological asset accounting, thus helping the spatial cognition and sustainable management of the state system in a more intuitive way[5,6]. However, due to the different parameter choices of different scholars, the evaluation results of the same ecosystem services may vary greatly, and there is a lack of comparability between the ESV obtained by different pricing methods, while the mature pricing method of ESV has not yet been formed internationally [7-10].

At present, the research on the evaluation method of ESV can be roughly divided into two categories: (1) A method based on the service function price per unit area. This method evaluates some key service functions by means of a series of ecological equations, such as food production, soil and water conservation, carbon and oxygen production, and habitat quality [11-14].The functional value method can accurately measure the size of some service functions in the region. However, for different service functions, different ecological equations and parameter inputs are often required, and the calculation process is more complicated [3]. Therefore, this method is mostly applied on a small scale, the implementation cost is large. In addition, when using this method for evaluation, scholars often lack the consideration of the ecological background of the study area. Moreover, there is no standard in choosing which service functions to be evaluated [15]. These drawbacks bring huge uncertainty to the evaluating results and limitations on results comparison. (2) A method based on value equivalent factor per unit area. This method was first proposed by Costanza et al [5]. This method divides different land ecosystems and service functions, obtaining the equivalent value based on meta-analysis and the area of each ecosystem to get the regional ESV. Compared with the functional value method, this method evaluate ESV more effectively on a large scale [16], and is widely used in research [3,5,17-19].

However, scholars have found that the evaluation results of the equivalent factor method are valid and reliable only when the equivalent factor accurately reflects the ecological background in the study area[16,20,21]. The equivalent factor proposed by Costanza et al.[5, 17] is aimed at global-scale value assessment, which is not consistent with the real ecological situation in China Xie et al.[18,19] conducted a questionnaire survey on Chinese ecologists, and put forward an equivalent factor table of ES for China in 2003 and 2008.In 2015, Xie et al.[3] updated and improved the equivalent factor table by combining various literatures and regional biomass, which is also the most scientific and systematic equivalent factor table in China. The equivalent factor table proposed by Xie et al.[3] essentially reflects the average level of the national ecosystem service function. At present, a large number of studies [14] showed that the strength of the different service function is affected by different ecological processes and conditions, for instance, organic matter production, gas regulation and nutrient cycling function [12] are closely related to net primary productivity (NPP); water supply and regulation function are closely related to rainfall soil erosion [23], habitat quality [13] and accessibility of recreational site [24]. Therefore, when the equivalent factor method is used to evaluate the ecological value of a region, corresponding spatial correction of equivalent factor is needed [8,25]. At present, scholars only use biomass or NPP to modify all types of service functions [3,18,19,26], which obviously does not match the real situation. Xie et al. [18] for the first time selected other ecological indicators (rainfall and soil retention) besides NPP to modify the service function.

Gansu Province is located in the northwestern inland in China. It is located at the intersection of the three major plateaus of the Loess Plateau, Qinghai-Tibet Plateau, Inner Mongolia Plateau and the arid regions of Northwest China, the Qinghai-Tibet Alpine Region, and the eastern monsoon region. Its special geographical location and natural conditions form a more distinct ecological structure. The types of ecosystems are complex and diverse in this region, covering forests, grasslands, deserts, wetlands, farmland, cities and other six continent ecosystems. The diversity of ecosystem types has caused significant regional differences. However, the current researches mostly focus on single or several ecosystems, and only investigate the several ecosystem service functions on the ESV in Gansu Province, such as forests[27-29], grassland[30,31], cultivated land[32]. Considering such complex ecological environment characteristics in Gansu Province, previous study has not investigate the ESV for the regional differences in space, and no value evaluation method has been established according with the ecological environment in the region.

This study takes into account the regional differences and the simplicity of the equivalent factor method. In view of the application of the equivalent factors of various ecosystems on a large scale, it is necessary to closely relate the equivalent factors with the national large scale and to the actual situation of Gansu Province. Based on a more refined classification of the ecosystem types in Gansu Province, the study added some ecosystem equivalent factors, constructed a revised index, and revised the equivalent factors studied by Xie et al.[3] to form an equivalent factor table that was suitable for assessment of ESV in Gansu Province. We constructed 11 regional differential adjustment indexes and readjusted the value of different service functions. Finally, we constructed a regional differential value evaluation model to evaluate the change of ESV in Gansu Province from 2000 to 2015. Considering the increasingly severe shortage and overuse of ecological services in region, the results of this study can provide scientific basis and decision support for local governments to make relatively complete ecological compensation policies.

## Materials and Methods

### The case study

The case study region is located in northwest China (Fig 1), at the intersection of three major plateaus—the Loess Plateau, the Qinghai Tibet Plateau, the Inner Mongolia Plateau—and three natural regions—the northwest arid region, the Qinghai Tibet alpine region and the eastern monsoon region. Gansu Province is a long and narrow region, covering a total land area of 425,800 km2, with complex and diverse geological landforms and climate types. In addition to the marine ecosystem, there are six main land use or cover types including forests, grasslands, deserts, wetlands, farmland, and urban areas. The Gansu region is part of the Chinese “two screens and three belts” strategic ecological security barriers, which aim to maintain and protect the survival and reproduction of organisms, maintain the natural ecological balance, and guarantee people’s livelihoods on the Qinghai Tibet plateau, the Sichuan-Yunnan Loess Plateau and the north sand belt. It is an important water conservation and supply area in the upper reaches of the Yangtze River and the Yellow River.

**Fig 1.**
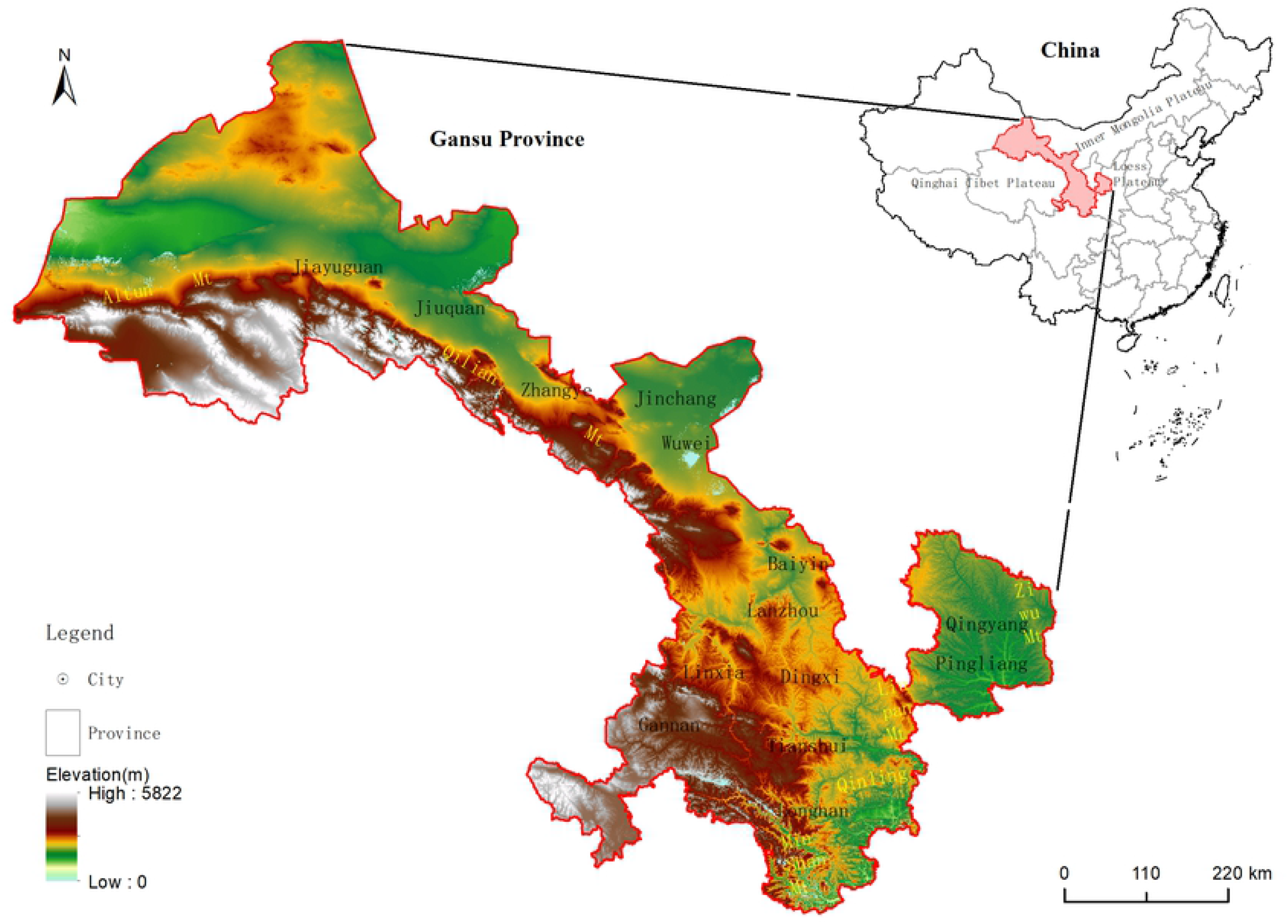
The location of the study area.

### Data source

#### The data of ecosystem type

The data of ecosystem types are from the satellite application center of the ministry of ecology and environment, and the period is 2000、2005、2010 and 2015. According to the actual situation of Gansu Province and the research needs, the ecosystem types are integrated into 7 primary types and 21 secondary types in the research area, and the corresponding database is established.

#### Meteorological data

In this study, the monthly average temperature, precipitation and sunshine hours are selected from 1981 to 2012 in Gansu Province and its surrounding meteorological stations. The data comes from Gansu Meteorological Bureau and China Meteorological science data sharing service network (http://cdc.nmic.cn).

#### Other geographic data

The annual average NPP data and the annual average water production data from 2000-2015 were used in this study, which comes from the satellite application center of the Ministry of ecological environment.

#### Socio-economic data

The social and economic data were used in this study from 2000-2015, and come from Gansu Province Statistical Yearbook, China Statistical Yearbook and national agricultural product cost-benefit data collection; the cultivated land quality data are from the annual renewal evaluation and monitoring results of cultivated land quality in Gansu Province (2017), and the grain output of each county is from Gansu Province Rural Yearbook (2000-2014); the monitoring data of atmospheric environmental quality status comes from Gansu environmental monitoring center station, with the period of 2015-2018. The monitoring data of surface water section comes from the bulletin of environmental conditions in Gansu Province and the bulletin of environmental conditions in China, with the period of 2000-2015.

### Classification of ecosystem service functions

On the basis of the research results of Costanza et al. [5], de Groot et al. [33], MA [34], Burkhard et al. [35] on the classification of ES and on the characteristics of the ecosystems in Gansu, ES were divided into: ecological integrity, regulatory services, supply services, and cultural services. Because of the problem of repeatability when measuring ES for ecological integrity, this was not included in the calculation of ES value. Supply services mainly considered crops, livestock and fresh water; regulation services mainly considered local climate regulation, air quality regulation, groundwater supply, soil conservation, windbreak and sand fixation, and water purification; and cultural services mainly considered entertainment and aesthetic value. Because Gansu is rich in biodiversity and this forms an important part of the value of ecological resources, the value of biodiversity protection was also included in the value accounting.

### Improvement method of value equivalent factor per unit area

#### Determination of standard equivalent factor

The research year is 2000-2015 in this study, the net profit of agricultural products is different in each year due to the different age, social and economic conditions and agricultural production technology. The ESV calculated only by the net profit of agricultural products in that year is not comparable. Therefore, according to the statistical information (for example, Gansu Yearbook in 2000-2015, data compilation of the second national agricultural census in Gansu Province, Gansu survey Yearbook, compilation of cost and income data of agricultural products in China), the sowing area and net profit of wheat, corn, potato and oil crops per unit area are obtained in Gansu Province from 2000 to 2015, and the average sowing proportion and average net profit are calculated. Based on this, the value of a standard equivalent factor is calculated. The calculation method is as follows:

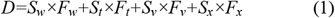

Where, *D* represents ESV of a standard equivalent factor (Yuan · hm-1); *S*_*W*_, *S*_*t*_, *S*_*V*_ and *S*_*X*_ represent sowing area proportion of wheat, corn, potato and oil to the sowing area of the four crops; *F*_*W*_, *F*_*T*_, *F*_*V*_ and *F*_*X*_ represent the average net profit of wheat, corn, potato and oil crops per unit area in Gansu Province (Yuan · hm-1).

#### Value equivalent factor per unit area

The basic value equivalent of ecosystem service function per unit area (hereinafter referred to as basic equivalent) refers to the annual average value equivalent of various service functions of different ecosystems types per unit area. Basic equivalent reflects the annual average value of different ecosystems and their various ecosystem service functions. Previous studies on equivalence factors [3,18,19] are based on the annual average value in national scale, and have a rough classification of ecosystem types, which cannot meet the needs of the refinement of ecosystem classification, nor more precisely reflect to the difference of service function among ecosystem types. Therefore, in this study, the average value equivalent factor per unit area of different ecosystems is determined in Gansu Province, with reference to the following calculation process.

1) For the types of ecosystem and the corresponding ecosystem service types in Gansu Province, if there is such kind of equivalence factor in the equivalence factor table of Xie et al. [3], this national average value equivalence factor will be used. On this basis, by constructing the correction coefficient, it will be converted into the average value equivalence factor of ecosystem service function in Gansu Province, such as the fresh water supply, local climate regulation, entertainment aesthetic value, air quality regulation, water purification and other ecosystem service function.

2) For the ecosystem types and corresponding ecosystem service types in Gansu Province, The equivalence factor table of Xie et al. [3] includes this category, but in this study, through the international literature system Elsevier, Springer, nature, Willey and the Chinese How Net database, search for the published relevant literature. We inputted the retrieval words of Gansu Province, the names of each basin and city of Gansu Province, Qilian Mountain, Gannan Plateau, etc. to collect and sort out the research results of this kind of ecosystem service value calculated by ecosystem service function quantity in Gansu Province. If there are many papers on the evaluation of ESV, the journals with high influence in the near future shall be selected for average calculation, and the proportion with standard equivalent shall be calculated as the basic equivalent of such ecosystem service function, such as local climate regulation and soil conservation function of shrub, etc.

3) According to the newly added ecosystem types and corresponding ecosystem service functions, we give priority to collect and sort out the domestic published research results of ecosystem service value calculated by ecosystem service function quantity in Gansu Province. The average of selected ESV should be calculated, then the proportion with standard equivalent also be calculated, so as to convert them into the average value equivalent factor of ecosystem service function in Gansu Province, as the basic equivalent of the ecosystem service function, which is used to determine the value equivalent of some ecosystem service functions, such as garden land, shrub land, forest land, swamp wetland, etc.

4) For the newly added ecosystem types and corresponding ecosystem service functions, if there are no relevant research results in Gansu Province, relevant research results in other regions in China shall be collected, ESV per unit area of ecosystem shall be calculated, then compared with standard equivalent value. It can be converted into average value equivalent factor of ecosystem service function in Gansu Province by constructing correction coefficient, as the basic equivalent of the ecosystem service function, which is used to determine the value equivalent of some ecosystem service functions, such as lakes, reservoirs, saline alkali land, urban green space, etc.

5) Considering the service functions of some ecosystems, there are great differences between Gansu Province and the whole country, although the equivalence factor table of Xie et al.[3] includes this types, but in this study, the value equivalence factor were localized and calibrated by calculating the ecosystem service function quantity per unit area, such as the biodiversity maintenance value of forest land, grassland, wetland and desert ecosystem, and the crop supply service of paddy field and dry land.

6) If there is no service function value of directly corresponding documents in the secondary classification of ecosystem, and it is not easy to calculate the ESV according to the existing documents, refer to the equivalent factors of Xie et al. [3] for the primary types are determined by experts’ experience, and at the same time, they are transformed into the average value equivalent factors of the ecosystem service function in Gansu Province through the construction of the correction coefficient, as the basic equivalent of the ecosystem service function, such as the local climate regulation and air quality regulation of the secondary types of desert and wetland ecosystem, and the soil conservation services and water purification services of second level of desert ecosystem.

Through the above six process, we get the value of main ecosystem types for a certain ecosystem service function per unit area in Gansu Province by referring to relevant literature or calculation, as shown in Table 1 below.

**Table 1.**
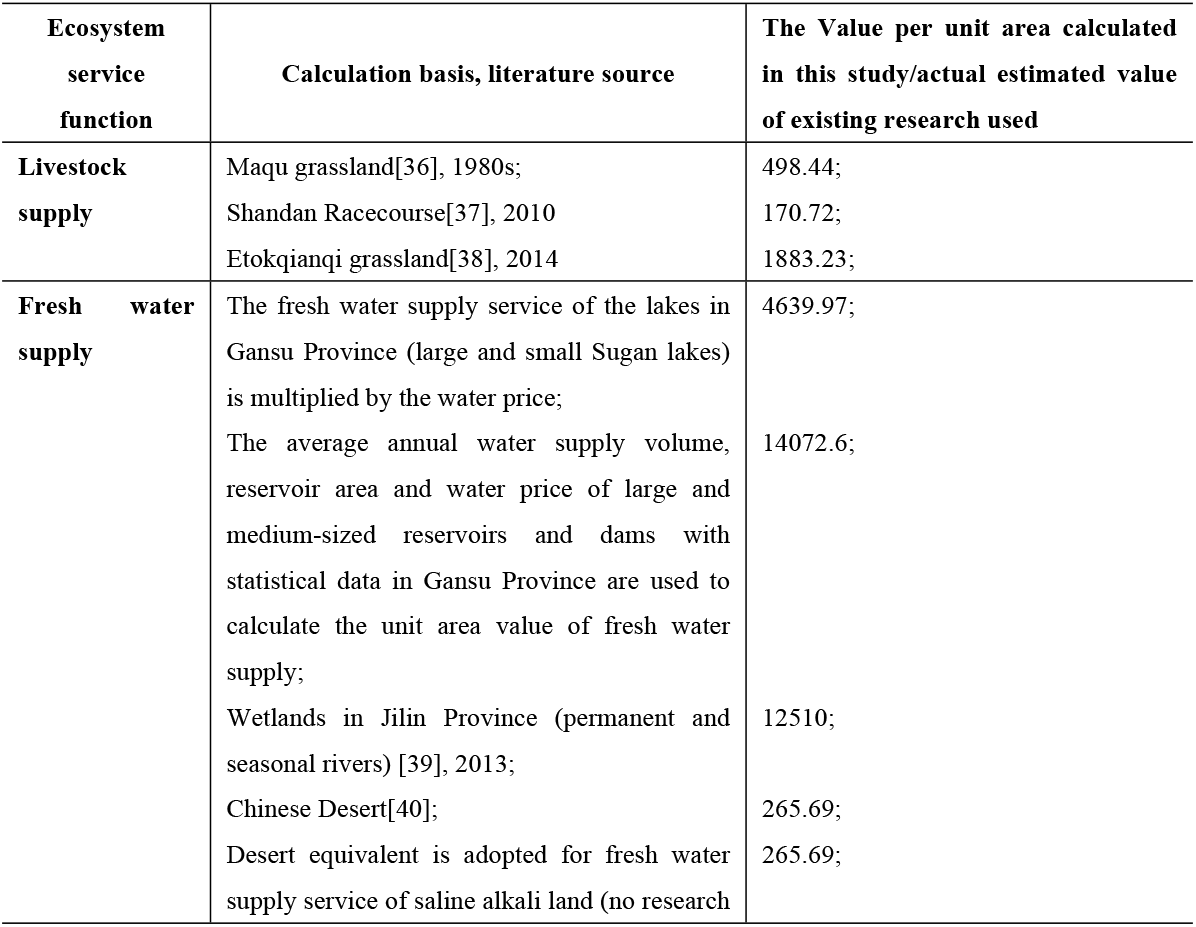
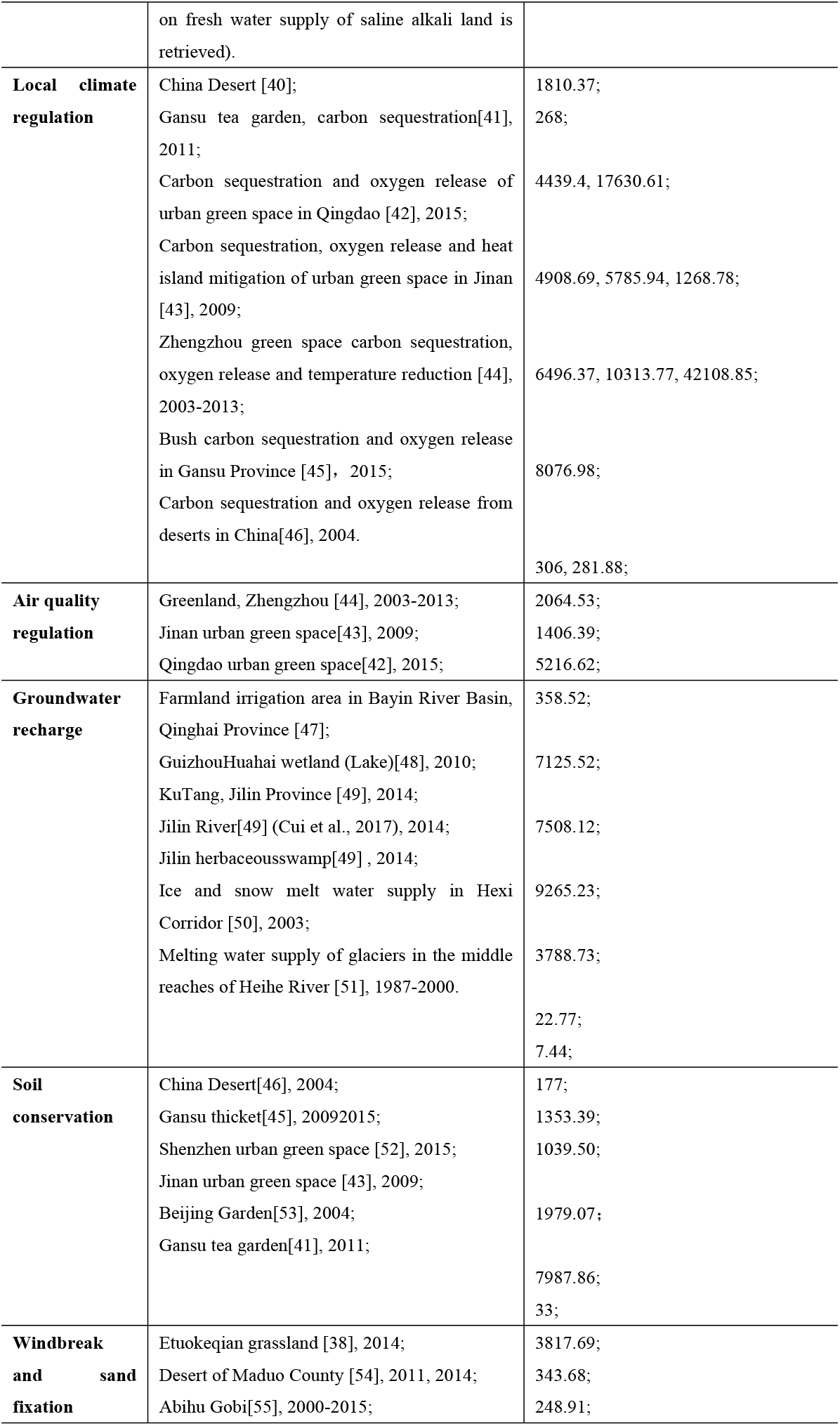
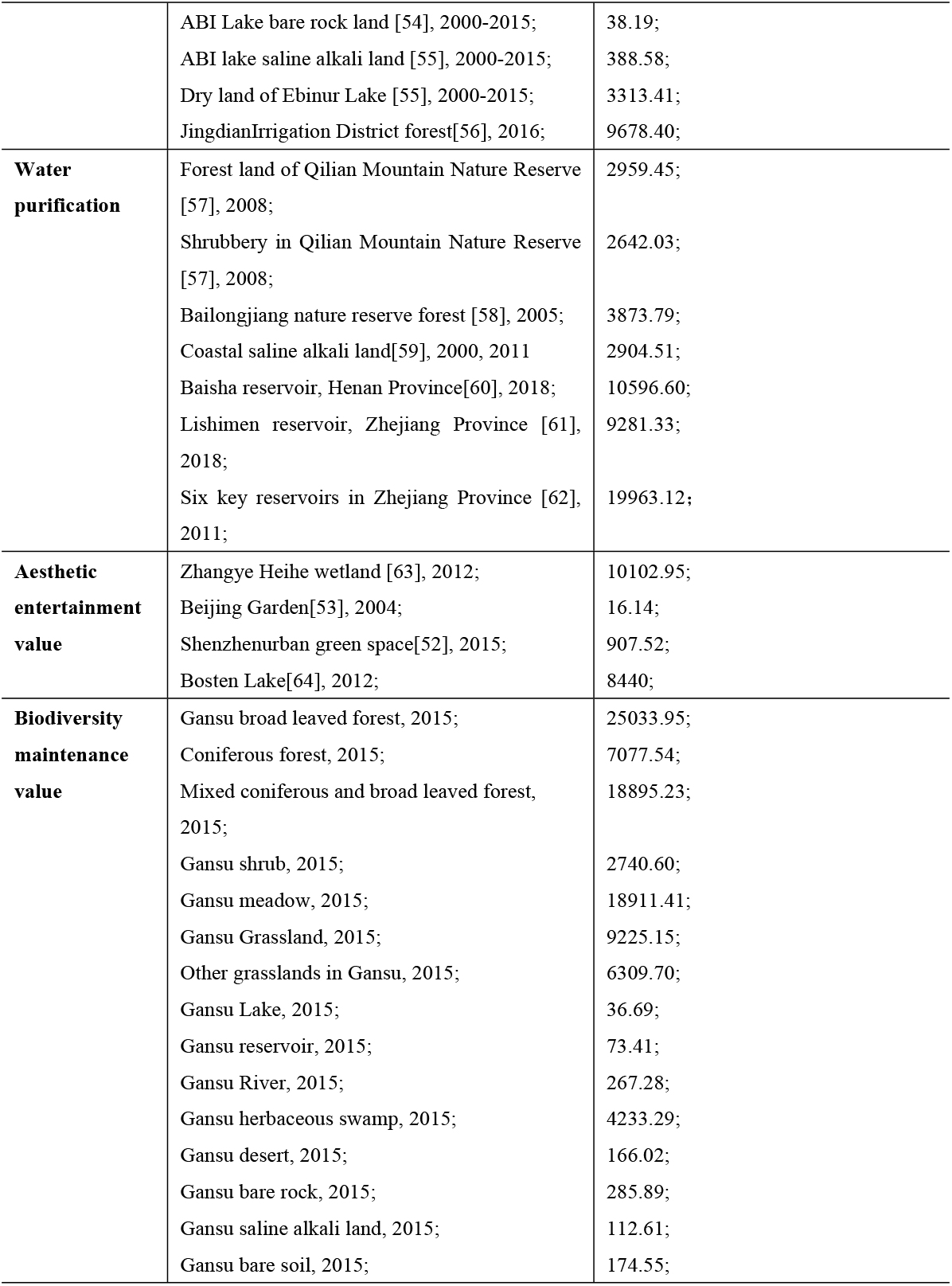
Calculation basis of value equivalent factor per unit area in Gansu Province.

#### Correction index of value equivalent factor per unit area

1) Crop supply correction index(*N*)

The calculation method of crop supply correction index is as follows:

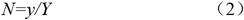

Where, *y* is the average output per unit area in Gansu Province, and *Y* is the national average output per unit area.

The calculation of y is based on the following formula:

*y* = (average yield per unit area of wheat/average yield per unit area of wheat) × sowing proportion of wheat + (average yield per unit area of corn/average yield per unit area of corn) × sowing proportion of corn + (average yield per unit area of potato/average yield per unit area of potato) × sowing proportion of potato + (average yield per unit area of oil/average yield per unit area of oil) × sowing of oil Proportion.

The calculation method of *Y* is the same as that of *y*.

2) Fresh water supply correction index(*D*)

The fresh water supply correction index is calculated as follows:

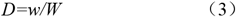

Where, *w* is the average water supply per unit area in Gansu Province (10000m3), and *W* is the average water supply per unit area in China (10000m3). The data of water supply comes from water resources bulletin (2000-2015) in Gansu Province and China.

3) Air quality regulation correction index (K)

The air quality regulation correction index is calculated as follows:

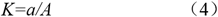

Where, *a* is the average proportion of air quality standards of prefecture level cities in Gansu Province, *A* is the average proportion of air quality standards of prefecture level cities in China, and the data of air quality standards comes from environmental quality bulletin (2000-2015) in Gansu Province and China.

4) Water purification correction index (*S*)

The calculation method of water purification correction index is as follows:

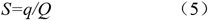

Where, *q* is the average length proportion of class I-III water reach in Gansu Province, and *Q* is the average length proportion of class I-III water reach in China. The length data of water quality reach is from water resources bulletin (2000-2015) in Gansu Province and China.

5) Entertainment aesthetic value correction index (Y)

The calculation method of entertainment aesthetics value is calculated as follows:

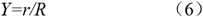

Where, *r* is the average tourism revenue per unit area in Gansu Province, *R* is the average tourism revenue per unit area in China, and the data of tourism revenue comes from Statistical Yearbook (2000-2015) in Gansu Province and China.

### Regional difference adjustment index

1) Crop supply regulation index (*A1*)

The calculation method of crop supply regulation index is as follows:

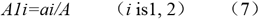

Where, *ai* is the average yield per unit area in Gansu Province, *A* is the average yield per unit area in Gansu Province, the calculation method of *ai* and *A* is the same as formula (1), *a1* is the high-yield area, *a2* is the low-yield area.

According to the research of Cheng [65], there are obvious spatial differences in the changes of grain production of cultivated land in Gansu Province. According to previous studies [66], the correlation between cultivated land quality and land use is relatively high, and the correlation coefficient reaches 0.874. Therefore, first of all, according to the cultivated land quality level (land use level) of each county and referring to related research [67] on the spatial distribution of land use level in 2015 in Gansu Province. Secondly, according to the proportion of cultivated land use level in the total cultivated land area of each county in Gansu Province, 27% of the counties higher than the medium proportion in Gansu Province are divided into high-yield areas, and the rest are divided into low-yield areas; Then according to the grain yield and sowing area of each county in high and low yield areas in Gansu Province, the average yield per unit area is calculated in high and low yield areas respectively, At the same time, calculate the average yield per unit area of the whole Province (refer to the calculation result of Formula 1). Compare the average yield in high-yield area and low-yield area with the average yield per unit area in Gansu Province, and get the regulation index of crop supply in high-yield area and low-yield area.

2) Livestock supply adjustment index(*A2*)

The livestock supply adjustment index is calculated as follows:

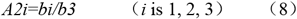

Where, *bi* is the average livestock carrying capacity of each area. According to relevant literature, *b1* is the average livestock carrying capacity in agricultural area, *b2* is the average livestock carrying capacity in semi pastoral area, and *b3* is the average livestock carrying capacity in pastoral area.

Gansu Province is divided into pastoral area, semi pastoral area and agricultural area according to the list of semi pastoral areas and counties in China because of the great difference of livestock supply capacity between pastoral area and agricultural area. According to relevant research [68], the livestock carrying capacity is calculated in the agricultural and pastoral areas in Gansu Province. The livestock carrying capacity in the agricultural areas is 0.85 times of that in the pastoral areas in Gansu Province. The average livestock carrying capacity in the agricultural and pastoral areas is taken as the livestock carrying capacity in the semi pastoral areas.

3) Fresh water supply regulation index(*A3*)

The calculation method of fresh water supply regulation index is as follows:

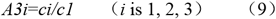

Where, *ci* is the average water yield per unit area of each area, *c1* is the average water yield per unit area in the high water area, *c2* is the average water yield per unit area in the poor water area, and *c3* is the average water yield per unit area in the dry area.

The distance between East and West, North and south is large in Gansu Province, and the precipitation decreases from southeast to northwest due to the influence of water vapor and terrain. According to the research [69], Gansu Province is divided into abundant water area (Liupanshan-Longshan area, Longnan mountain area, Gannan plateau, Qilian Mountain Area), poor water area (Longdong, Longdong Loess Plateau area, north of Lanzhou area) and dry area (Hexi Corridor, Beishan Mountain Area and its desert area bounded by Qilian mountain foot). Using the water yield module in invest model, the average multi-year water yield of high water area, low water area and dry area is calculated in 2000-2015 in Gansu Province. Finally, the water yield per unit area in the low water area is 0.68 times of that in the high water area, and the water yield per unit area in the dry area is 0.08 times of that in the high water area.

4) Local climate regulation index(A4)

The calculation method of local climate regulation index is as follows:

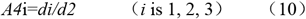

Where, *di* is the average NPP of each partition, *d1* is the NPP value of high adjustment area, *d2* is the mean value of NPP in the middle regulation area, and *d3* is the NPP value of low regulation area.

A large number of observation data analysis shows that, as the underlying surface of the climate system, the change of surface vegetation may have a significant impact on local and regional climate by changing surface attributes such as surface albedo, roughness, soil moisture [70-73]. The higher the area of vegetation net primary productivity (NPP), the stronger the function of climate adjustment. Therefore, the value of NPP is used to measure the regional difference of climate regulation. According to the spatial distribution characteristics of NPP and the boundary of township, Gansu Province is divided into high regulation area, middle regulation area and low regulation area. The average value of NPP in the three districts was calculated respectively, and the ratio of NPP was taken in the high regulation area and the middle adjustment area as the adjustment index in the high adjustment area. The ratio of the average value of NPP in the low adjustment area and the median adjustment area was used as the adjustment index of the low value area.

5) Air quality regulation index (*A5*)

The air quality regulation index is calculated as follows:

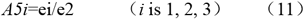

Where, *ei* is the average vegetation coverage in each area, *e1* is the average vegetation coverage in the area with good air quality, *e2* is the average vegetation coverage in the area with general air quality, and *e3* is the average vegetation coverage in the area with poor air quality.

Generally speaking, the better the air quality in a region, the greater the air quality regulation service function. According to 2015 environmental quality bulletin in Gansu Province, PM_10_ and PM_2.5_ are the main air pollutants; only one of the 14 cities and prefectures has reached the secondary standard of ambient air quality, so the concentration of the pollutant is taken as the index to measure the level of air quality regulation function. In this paper, PM_10_ and PM_2.5_ concentration monitoring data are selected in 111 provincial monitoring points in Gansu Province, and through Kriging interpolation, the spatial distribution of PM_10_ and PM_2.5_ concentration is obtained in the whole province, which is divided into three zones. The area meeting the secondary quality standard is classified as the area with the best air quality, indicating that the area has the highest air quality regulation function, and other areas are divided into two areas according to the concentration. According to the relevant research, vegetation coverage is closely related to the air purification function. In this paper, the ratio of the average vegetation coverage in three regions is used as the air quality regulation index.

6) Groundwater recharge regulation index (*A6*)

The calculation method of groundwater recharge index is as follows:

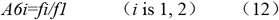

Where, *fi* is the ratio of actual exploitation amount and exploitable amount of groundwater in each zone, *f1* is the ratio of actual exploitation amount and exploitable amount of groundwater in over mining area, *f2* is the ratio of actual exploitation amount and exploitable amount of groundwater in non over mining area, assuming that the actual exploitation amount and exploitable amount of groundwater in non over mining area are in balance, the ratio is set as 1.

The overexploitation of groundwater results in the drainage of aquifer, the decrease of groundwater level, the formation of funnel, and land subsidence. Therefore, when the rapid development exceeds the resource stock and environmental capacity, the value of groundwater ecosystem will inevitably continue to appreciate. Different regions have different needs for groundwater recharge function, resulting in different values. In order to reflect the regional difference of groundwater recharge regulation function value, we uses groundwater over mining area and non over mining area to measure the regional difference of groundwater recharge function. Groundwater over mining area has higher groundwater recharge value than non over mining area. According to the delimitation results of groundwater over mining area in Gansu Province [74], there are 46 over mining areas, involving 32 counties. The ratio of the actual and exploitable groundwater in the over mining area is used as the groundwater supply regulation index in the over mining area.

7) Soil conservation regulation index (*A7*)

The calculation method of soil conservation regulation index is as follows:

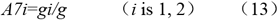

Where, *gi* is the average erosion modulus of each area, *g1* is the average erosion modulus of key prevention area, *g2* is the average erosion modulus of key control area, and *g* is the allowable amount of soil erosion.

Gansu Province is located at the junction of three plateaus, and its soil conservation functions vary greatly in different areas. According to distribution range of soil and water loss in the key prevention areas and key control areas designated in Gansu Province, there have better vegetation, less soil and water loss, stronger soil conservation functions in the key prevention areas, but have low forest and grass coverage, fragile ecological environment, and serious soil and water loss in the key prevention area. Therefore, the whole Province is divided into two zones according to the range of the key prevention and control areas. At the same time, according to the classification standards of soil erosion, the allowable amount of soil and water loss in the Northwest Loess Plateau is 1000t/km2. Based on the ratio of the average erosion modulus and the allowable amount of soil and water loss in the two zones, the adjustment index of soil conservation is constructed.

8) Regulation index of windbreak and sand fixation (*A8*)

The calculation method of windbreak and sand fixation regulation index is as follows:

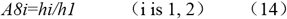

Where, *hi* is the amount of windbreak and sand fixation in each zone, *h1* is the amount of windbreak and sand fixation in the service area of windbreak and sand fixation, and *h2* is the amount of windbreak and sand fixation in other areas.

The Hexi Corridor in the north of Gansu Province and the surrounding county of Qingyang City is located in the desert Gebi area. Therefore, this area is classified as a service area for windbreak and sand fixation, while other areas are not considered to have the function of windbreak and sand fixation.

9) Water purification regulation index (*A*9)

The calculation method of water purification regulation index is as follows:

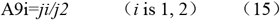

Where, *ji* is the target proportion of water quality in each zone; *j1* is the length proportion of class II and above water reaches in high water purification area; *j2* is the length proportion of class III and below water reaches in low water purification area.

Xie et al. [75] pointed out that as the pollution of rivers and lakes is becoming more and more serious, the water quality regulation function of rivers is becoming lower and lower, and rivers and lakes almost become the place for waste. Generally speaking, the water quality of the reach is closely related to its water purification function. If the water quality of this area is significantly better than that of other areas, the water purification function of this area is of great importance. In this study, the water quality objectives of 236 monitoring sections of the river in Gansu Province are used to measure the water purification function of the region where the river is located, and the water quality objectives of each county are counted. If the water quality objectives of class I, class II or class II water and class III water reaches can be achieved simultaneously, the water purification function of the county is considered to be high, and that of other water quality counties is considered to be water purification function low. At the same time, the length proportion of class II water reach and class III water reach and below are calculated. The length proportion of class II water reach to class III water reach and below is taken as the water purification regulation index in high water purification area.

10) Entertainment aesthetics value adjustment index (*A10*)

The value adjustment index of entertainment aesthetics is calculated as follows:

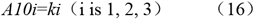

Where, *ki* is the value adjustment index of entertainment aesthetics in each zone, *k1* is the value adjustment index of entertainment aesthetics in key tourist areas, *k2* is the value adjustment index of entertainment aesthetics in general tourist areas, and *k3* is the value adjustment index of entertainment aesthetics in other regions.

Entertainment value refers to the value obtained by tourists when they are engaged in tourism activities in the ecotourism scenic spot, which is the sum of the use value brought by direct recreation and the non use value possessed by resources; aesthetic value refers to the pleasure value brought by natural ecosystem to people’s aesthetic perception by its natural landscape and cultural landscape directly bred, and the value of its own objective aesthetic attribute belongs to non use value. If people cannot reach the area that can bring recreational and pleasure value to people, it is considered that the area cannot provide the service function or temporarily does not have the function. Based on this, we first determine that the core area and buffer area of the Nature Reserve cannot provide this service temporarily. Other nature protected place such as Forest Park, Geopark, Scenic Area, Wetland Park, etc. are the key tourism areas, which provide the highest entertainment aesthetic value. Secondly, the whole tourism county (excluding the key tourism areas) is regarded as the general tourism area, and other areas enjoy the entertainment aesthetics value is the lowest. The adjustment indexes of different regions are assigned by expert judgment.

11) Biodiversity maintenance value regulation index (*a11*)

The value adjustment index of biodiversity maintenance is calculated as follows:

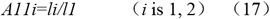

Where, *li* is the habitat quality index of each region, *l*_*1*_ is the habitat quality index of priority area for biodiversity conservation; *l*_*2*_ is the habitat quality index of other regions.

According to the conservation plan of biodiversity priority area in Gansu Province, there are seven biodiversity priority areas in Gansu Province. This paper considers that the areas located in the priority areas have the highest biodiversity maintenance value, followed by other areas, so the province is divided into two areas. In order to determine the adjustment index of different regions, this paper uses the habitat quality module of invest model to calculate the habitat quality index of different regions, and determines the adjustment index by comparing the size of the habitat quality index in the two regions.

Through the above methods, the value equivalent factors table per unit area is established in Gansu Province (Table 2).

**Table 2.**
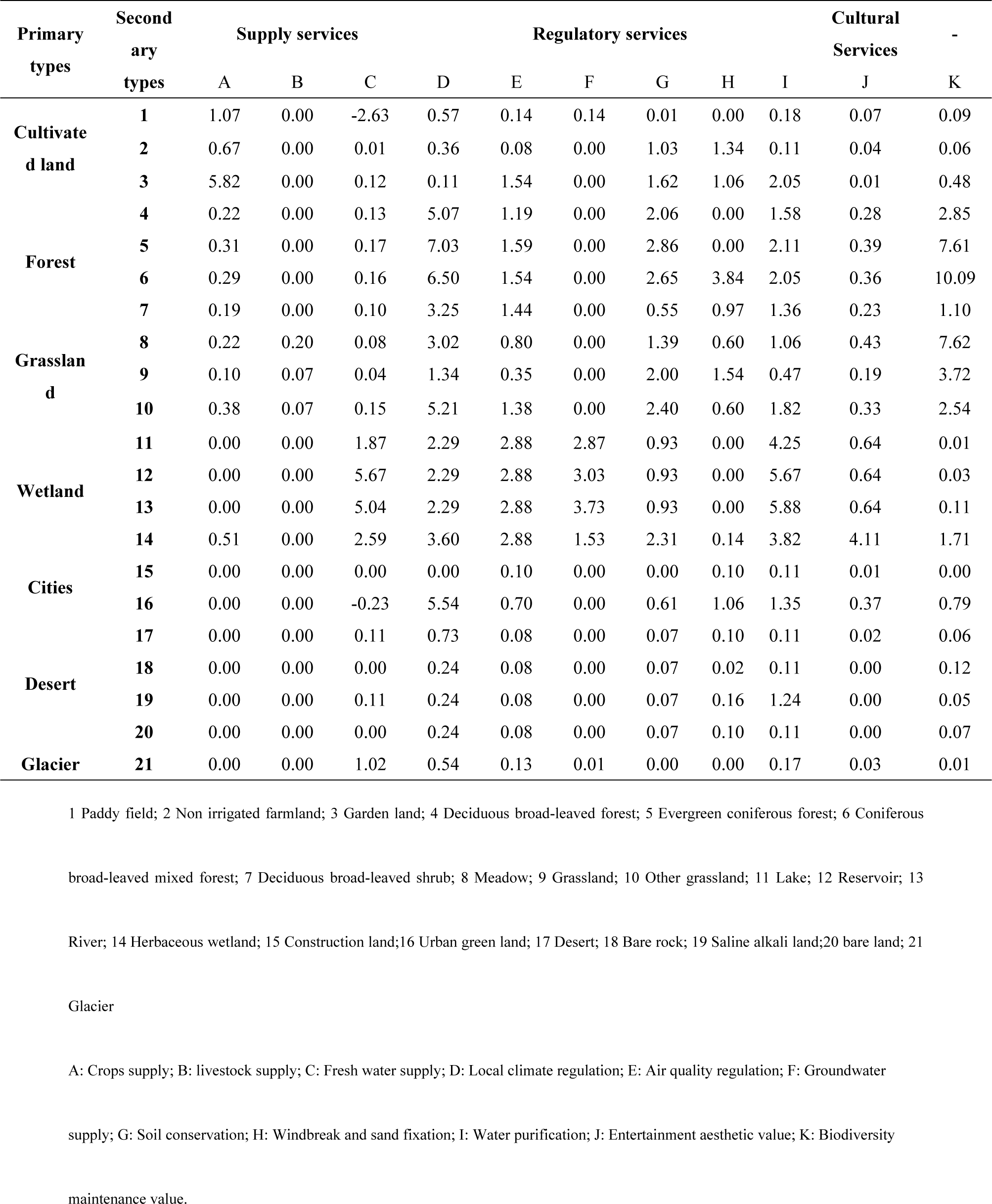
The value equivalent factors per unit area in Gansu Province.

### Value evaluation model based on regional difference

Different geomorphic types will affect the distribution of light, heat, water and soil types in the region [76]. Different regional ecological environment, degree of ecological protection, intensity of ecological demand for different land use types and implementation of local policies will affect the benefits human beings get from the ecosystem, thus affecting the regional differences and divisions of ESV. The ArcGIS spatial analysis tool is used to grid Gansu Province, and a complete grid of 1km×1km is extracted. Based on the calculation of ESV of grid unit, the total ESV is calculated in Gansu Province by the following formula (Fig 2). The total ESV in Gansu Province (*V*) can be expressed as:

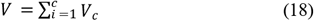

**Fig 2.**
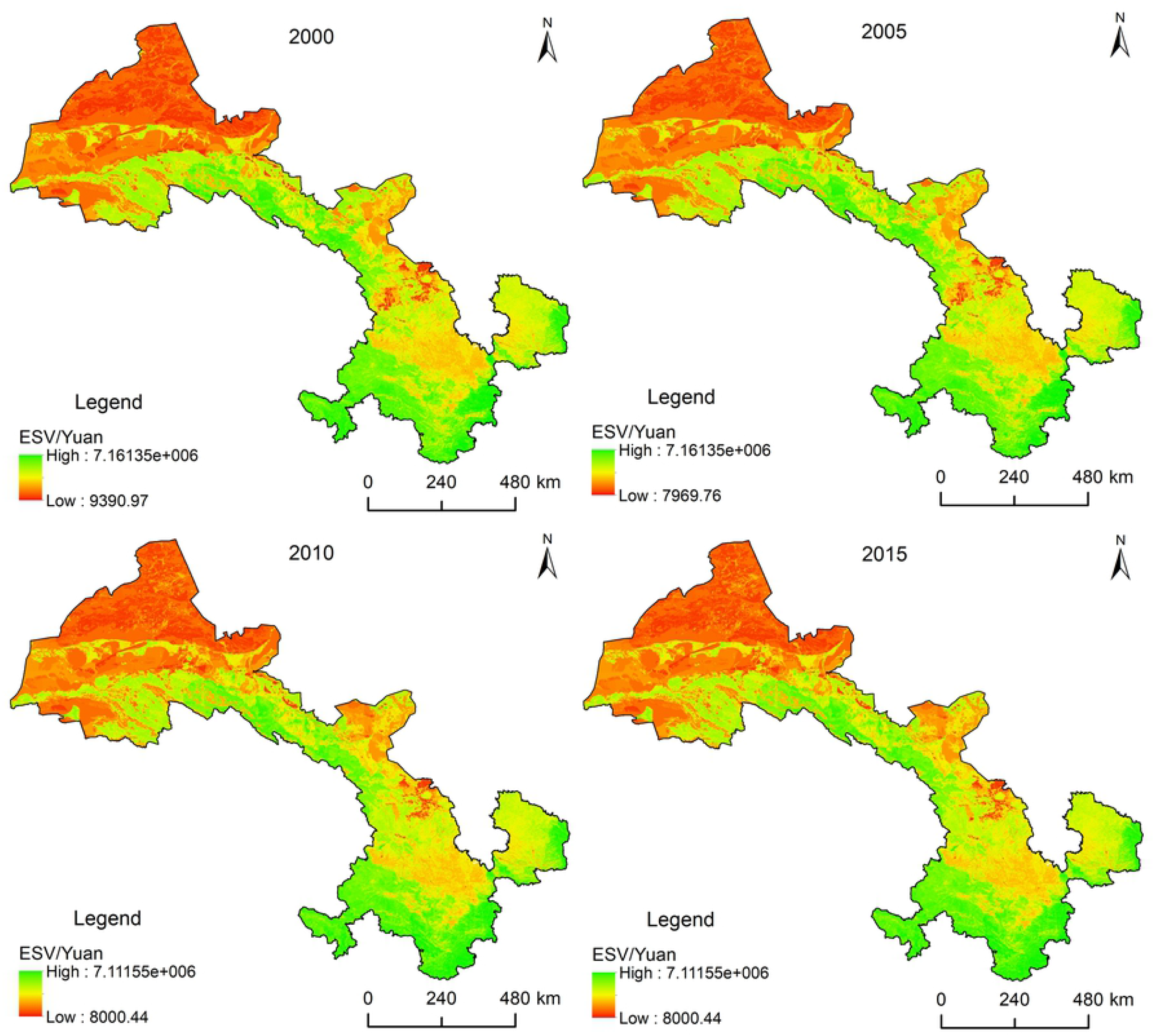
The flow chart of ecosystem service value evaluation model based on regional differences.

Where, *V*_*c*_ is the value of ecosystem service function *c, c* is ecosystem service function, and the value is 1, 2…. 11.

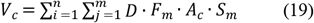

Where, *D* is the standard equivalent factor, *F*_*m*_ is the value equivalent factor per unit area of ecosystem type m, *A*_*c*_ is the regional difference adjustment index of ecosystem service function *c, S*_*m*_ is the area of ecosystem type m (ha), *n* is the grid number.

## Results

### Analysis on the change of ecosystem in Gansu Province

From the composition of ecosystem types in Gansu Province (Table 3), desert is the largest type in Gansu Province, followed by grassland, arable land, forest ecosystem. Both glacier and wetland ecosystem only account for a small part. Desert are mainly distributed in the north of Gansu; grasslands are mainly distributed in the Gannan plateau, central and Eastern of Gansu; cultivated land is mainly distributed in the middle of Gansu, Hexi Corridor; and forests are mainly distributed in Longnan, Ziwuling and Qilian Mountains. According to the change of ecosystem area in Gansu Province, the forest, grassland and urban ecosystem have been increasing continuously in the past 15 years in Gansu Province. The cultivated land ecosystem has been decreasing continuously. Although the wetland and glacier ecosystem have been increasing in the past 15 years, they have been decreasing in general. The desert ecosystem has been increasing in general.

**Table 3.** Change of ecosystem pattern from 2000 to 2015 in Gansu Province (km2)

| Types of ecosystem | 2000 | 2005 | 2010 | 2015 |
| --- | --- | --- | --- | --- |
| Forest | 54372.71 | 55118.01 | 56178.19 | 56228.06 |
| Grassland | 120419.34 | 122061.74 | 124629.60 | 124739.27 |
| Wetland | 2624.82 | 2732.48 | 2440.76 | 2545.59 |
| Cultivated land | 76454.39 | 74362.82 | 68743.14 | 68328.45 |
| Cities | 3438.61 | 3723.99 | 4071.09 | 4624.28 |
| Desert | 167223.90 | 166546.96 | 168485.42 | 168076.37 |
| Glacier | 909.06 | 896.83 | 894.63 | 900.81 |

### Overall evaluation of ESV in Gansu Province

In general, ESV decreases from south to North and from east to west in Gansu Province (Fig 3), which is consistent with the spatial distribution of forest, grassland and desert ecosystem. There are contiguous desert areas in the north of Gansu Province with the lowest ESV. However, the ecological environment is relatively good in the south of Gansu Province, with high vegetation coverage and ESV. In the Middle East of Gansu Province where human activities are frequent, which induced relatively high disturbance on the natural ecological environment, thus ESV is relatively low.

**Fig 3.**
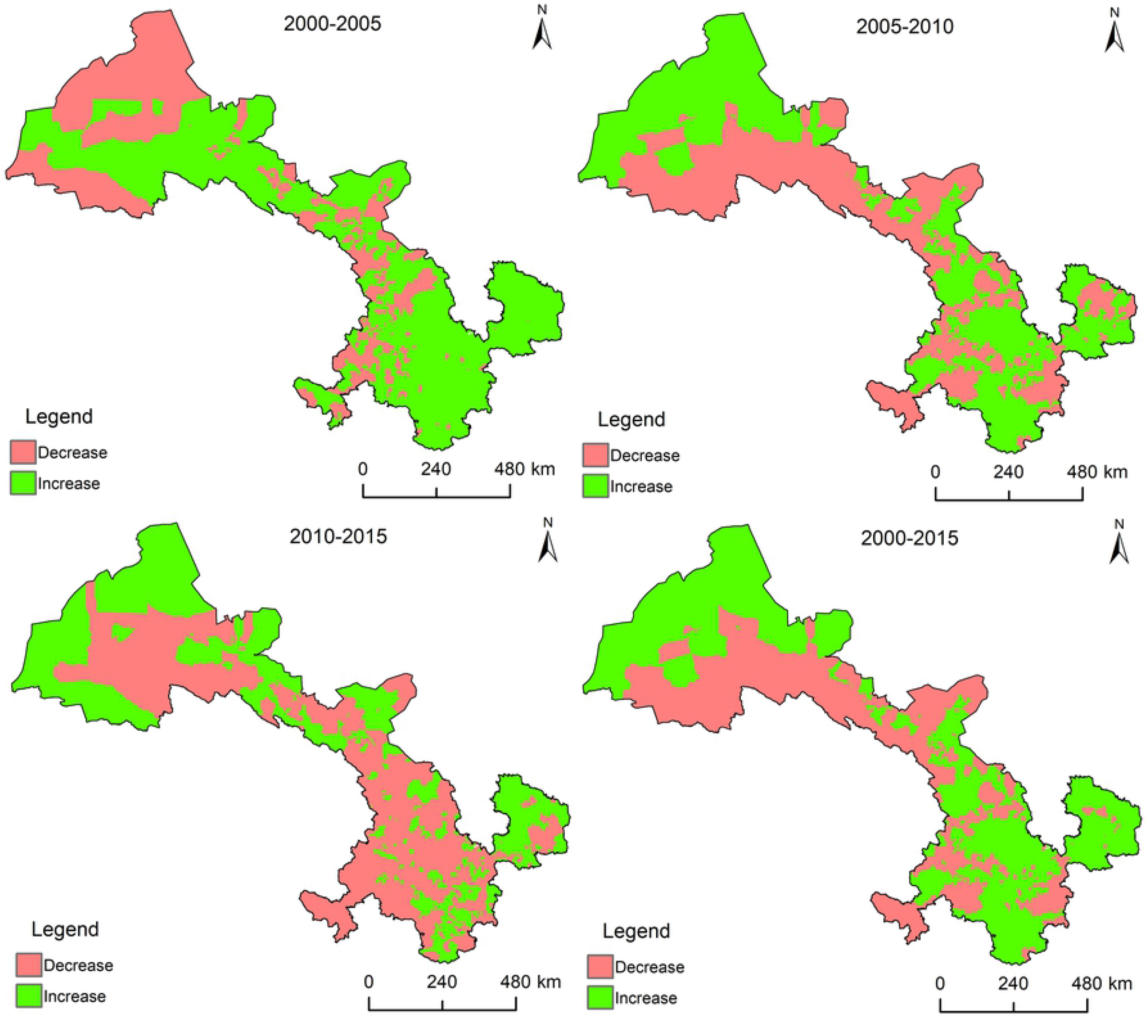
The ESV from 2000 to 2015 in Gansu Province.

From the value of each service function of ecosystem (Table 4), the value of local climate regulation and biodiversity maintenance is much higher than that of other service functions, which constitutes the main component of ESV in Gansu Province. From the change of each service function value, the value of soil conservation, windbreak and sand fixation, biodiversity protection has been increasing continuously in the past 15 years, while the supply value of crops is decreasing continuously. The supply value of fresh water, local climate regulation, air quality regulation, water purification function, groundwater supply and entertainment aesthetic value show a fluctuating change state and a decreasing trend as a whole.

**Table 4.** The ESV from 2000 to 2015 in Gansu Province (108Yuan)

| ES | 2000 | 2005 | 2010 | 2015 |
| --- | --- | --- | --- | --- |
| Crops supply | 965.67 | 958.54 | 886.63 | 879.29 |
| Livestock supply | 92.81 | 93.78 | 93.03 | 93.09 |
| Fresh water supply | 199.34 | 201.60 | 197.55 | 198.90 |
| Local climate regulation | 8216.79 | 8301.09 | 8164.14 | 8171.22 |
| Air quality regulation | 1961.98 | 1978.94 | 1925.38 | 1928.47 |
| Groundwater recharge | 70.64 | 76.43 | 57.67 | 63.62 |
| Soil conservation | 1551.19 | 1559.10 | 1605.89 | 1606.29 |
| Windbreak and sand fixation | 728.68 | 734.78 | 772.64 | 774.98 |
| Water purification | 2932.81 | 2958.28 | 2819.21 | 2824.91 |
| Aesthetic value of entertainment | 425.42 | 427.25 | 418.56 | 418.76 |
| Biodiversity maintenance value | 5215.96 | 5278.48 | 5428.37 | 5436.01 |
| Total value | 22361.29 | 22568.27 | 22369.07 | 22395.55 |

From the value composition of each ecosystem type (Table 5), the value of grassland and forest ecosystem is the highest, accounting for more than 75% of the total value, while the value of urban and glacial ecosystem is lowest. From the change of each ecosystem type value, the value of forest and urban ecosystem is increasing, while the value of cultivated land ecosystem is decreasing; The value of grassland, wetland and glacier ecosystem is fluctuating, and the overall trend is decreasing. In contrast, the value of desert ecosystem is fluctuating, and the overall trend is increasing. Over the past 15 years, the total value of various ecosystem services has increased by 3.426 billion Yuan, and the increase of forest ecosystem value is the most, while the decrease of grassland ecosystem value is the most.

**Table 5.** Value of each ecosystem type from 2000 to 2015 in Gansu Province (10^8^Yuan) and proportion (%)

| Ecosystem | 2000 |  | 2005 |  | 2010 |  | 2015 |  |
| --- | --- | --- | --- | --- | --- | --- | --- | --- |
|  | Value | Proportion | Value | Proportion | Value | Proportion | Value | Proportion |
| Forest | 7193.20 | 32.17 | 7312.00 | 32.40 | 7477.01 | 33.43 | 7489.26 | 33.44 |
| Grassland | 10222.26 | 45.71 | 10352.27 | 45.87 | 10181.71 | 45.52 | 10188.37 | 45.49 |
| Wetland | 371.43 | 1.66 | 383.54 | 1.70 | 348.94 | 1.56 | 359.66 | 1.61 |
| Cultivated land | 2969.03 | 13.28 | 2918.64 | 12.93 | 2671.66 | 11.94 | 2659.78 | 11.88 |
| Cities | 109.32 | 0.49 | 118.56 | 0.53 | 131.50 | 0.59 | 140.64 | 0.63 |
| Desert | 1484.10 | 6.64 | 1471.39 | 6.52 | 1547.69 | 6.92 | 1547.25 | 6.91 |
| Glacial | 11.95 | 0.05 | 11.86 | 0.05 | 10.55 | 0.05 | 10.59 | 0.05 |
| Total | 22361.29 | 100.00 | 22568.27 | 100.00 | 22369.07 | 100.00 | 22395.55 | 100.00 |

### Analysis on the spatio-temporal change of ESV in Gansu Province

From the spatial changes of ESV(Fig 4), over the past 15 years, the township with increased ESV are mainly distributed in the east of Gansu Province, the south of Gansu Province and the west of Hexi Corridor; the township with decreased ESV are distributed in Qilian Mountain and Gannan plateau.

**Fig 4.**
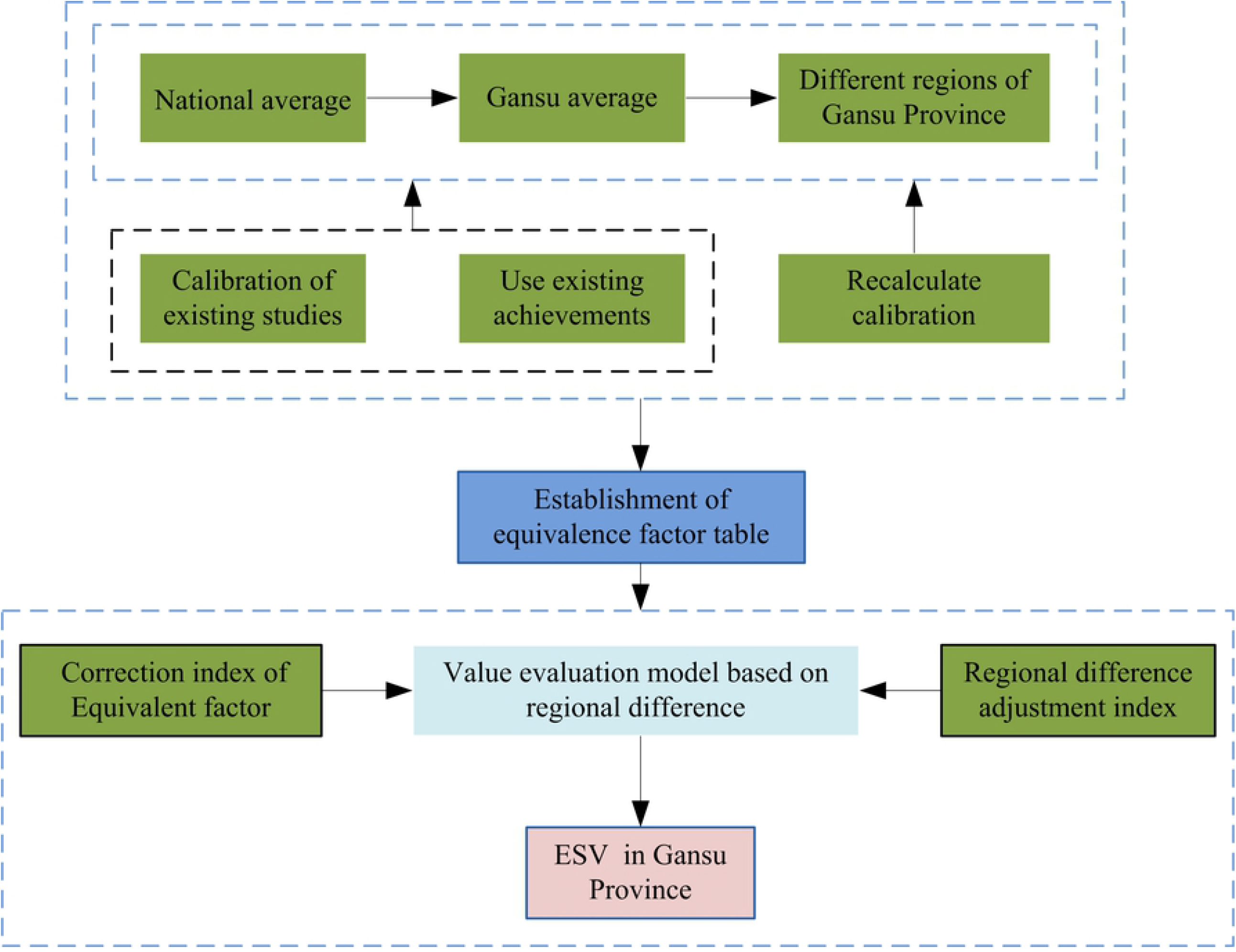
The change distribution map of ESV for each town from 2000-2015 in Gansu Province.

From the change of ecosystem service value every five years, over the past 15 years, the number of township with increased ESV has become less and less. There were 959 township increased in 2000-2005, 758 township in 2005-2010, and only 391 township increased in value in 2010-2015. From 2000 to 2015, there were 818 townships with increased ESV.

From 2000 to 2005, the township with increased ESV are mainly distributed in Qilian Mountains, Eastern and southern of Gansu (Tianshui, Pingliang, Qingyang and Longnan). From 2005 to 2010, ESV of most townships has decreased in Gannan plateau and Qilian Mountain, while the increased township were mainly located in Tianshui, Longnan and the western section of Hexi corridor. From 2010 to 2015, ESV of most township was declining in Lanzhou, Baiyin, Dingxi, Gannan plateau and Hexi corridor, and the value-added township were compressed to the north and south.

## Discussion

### Advanced Value Evaluation Model

In this study, ESV was evaluated by improving value equivalent factor per unit area and constructing a regional differential value assessment model in Gansu Province. Different ecosystem types provide different ES types to humans. In the current research on ESV, scholars mostly refer to the classification of ecosystem types by Costanza et al. [5], which is simply divided into forest, grassland, farmland, wetland, desert and river. Although Xie et al. [3] improved this method by accounting of the value of 14 types of ecosystem services in China, and more precisely reflected the differences between ecosystem types. However, this method is based on the national scale, which cannot meet the needs of practical research i.e. reduction of the research scale and refinement of the classification of ecosystems. According to the actual ecological situation in Gansu Province, this study identify 7 types of primary ecosystems and 21 types of secondary ecosystems to cover the main ecosystems types in Gansu Province more comprehensively, which reflect the differences between the types of ecosystems, and then accurately reflect the importance of ESV.

In different regions, even if the same ecosystem type provides different ES and their value to humans are quite different. Therefore, the value equivalent factor of Costanza et al.[5] and Xie et al. [3] is improved by scholars in practical research, and factors are introduced that could reflect regional differences, such as NPP, biomass, vegetation coverage, and soil erosion. However, these factors can only reflect the regional differences in the main types of ES functions that are mainly driven by natural factors. It is even more difficult to truly reflect differences of ecological service functions in the region with complex ecological environment characteristics and socioeconomic conditions in Gansu Province. In this study, firstly, correction index of value equivalent factor per unit area s constructed. We directly convert the equivalent factor on the national average level of Xie et al. [3] to the average level in Gansu Province; and directly calculate the average level of some equivalent factors in Gansu Province, and then convert the equivalent factors in the average level in Gansu Province to the level with significant regional differences by constructing the regional difference adjustment index of each ES function type. In the process of regional difference conversion, the differences have been fully considered in the ecological environment quality, resource endowment, and economic and social development of different regions. These can better reflect the ecological well-being of residents in different regions or the degree of damage to the local ecological environment and resources.

### Errors in the value evaluation model

In the value evaluation method based on the equivalent factor, the accurate construction of the equivalent factor table is the core of the equivalent factor method. In the process of improving the equivalent factor table of Xie et al. [3], this study integrated evaluation results from literatures based on the physical quantity method in Gansu Province or other regions in China. This can avoid or reduce the subjective conjecture which is easily caused by the empiricism from the past. The accuracy of the research results in the literature on different ES types will affect the size of the equivalent factor in this study. Considering the incomplete research results of different ecosystems or ecological service functions in Gansu Province, the ecological service functions of some ecosystem types are investigated based on the relevant research results from the available regions in China, which will inevitably lead to a certain amount of uncertainty in this study. Future research need to quantitatively calculate the physical quantity for the ecosystem type or ecological service function that are not available in current research results, and then to determine the equivalent factor to improve the relevant results of this research.

### Reliability of value evaluation models

ESV assessment research has always been a hot topic in other regions in China, while there are few related researches on ESV in Gansu Province. The existing research only focus on the value of a single ecosystem service on the provincial scale [27,32,77], and the calculation method of unit area value is the main method, however, the research on the provincial integrated ESV is almost blank at present[78]. The value of forest ecosystem services is747.70 billion Yuan in this study. This study does not consider the difference of people’s willingness to pay and ability to pay for ESV caused by the social development. By including the willingness to pay and ability to pay, the value of forest ecosystem services is 1971.228 billion Yuan in 2010, which is the same with the official release of service value of forest ecosystem (2007.97 billion Yuan in 2011), and the service value of forest ecosystem assessed by Wang et al. [27] (2163.86 billion Yuan). However, there is a big difference between the ESV evaluated in Gansu Province by Qi [78] and the ESV in this study. The value per unit area is only 98.569 billion Yuan in 2010 [78] by using the value calculation method per unit area. Obviously, the assessment result is significantly smaller.

## Conclusions

Based on the characteristics of ecosystem in Gansu Province, this paper built up a revised index based on the increase of some ecosystem equivalent factors, and revised the equivalent factors studied by Xie et al. [3] to form an equivalent factor table in line with ecosystem service valuation in Gansu Province, then constructed 11 regional differences adjustment index to readjust the value of different service functions. The regional difference assessment model constructed in this paper distinguished the regional differences in the similar ecosystems services to evaluate the ESV in Gansu Province more objectively. The main conclusions are as follows:

(1) The desert ecosystem is the type with the largest area in Gansu Province, followed by grassland, arable land, and forest ecosystems, and the remaining ecosystems that account for only a small part. Desert mostly locates in northern Gansu. Grasslands are mainly distributed in the Gannan Plateau, Longzhong, and Longdong. Cultivated land is mainly distributed in Longzhong and Hexi corridors, and forests are mainly distributed in Longnan, Ziwuling, and Qilian Mountains.

(2) From 2000 to 2015, the grassland ecosystem area increased the most, while the cultivated land ecosystem area decreased the most. Forest, grassland, and urban ecosystems in Gansu Province continue increasing. Cultivated land ecosystems continue decreasing. Although wetlands and glacial ecosystems have increased during the past 15 years, they have generally decreased. Desert ecosystems have increased.

(3) In 2015, the total ESV reached 2239.555 billion Yuan in Gansu Province, among which forest and grassland ecosystems contribute the most. In terms of the value of each service function of the ecosystem, the local climate regulation and biodiversity maintenance functions are the main service functions in Gansu Province. On the spatial distribution of service values, ESV gradually increases from northeast to southwest, and the areas with high ESV concentration are Qilian Mountain and Longnan Mountain.

(4) From 2000 to 2015, ESV increased by 3.426 billion Yuan in Gansu Province. The value of forest ecosystems increases the most, while the value of grassland ecosystems decreases the most, showing a trend of increasing first, then decreasing, then slowly increasing. The value of forest and urban ecosystems continues to increase, and the value of cultivated land ecosystems continues to decrease. From the spatial characteristics of ESV changes, areas with reduced value gradually move from central Gansu to surrounding areas.

## Acknowledgments

We are also grateful for the comments and criticisms of an early version of this manuscript by our colleagues and the journal’s reviewers. Acknowledgement for the data support from “National Earth System Science Data Center, National Science & Technology Infrastructure of China. (http://www.geodata.cn)”

## References

1. Lu CX, Xie GD, Xiao Y, Yu YJ. Ecosystem diversity and economic valuation of Qinghai-Tibet Plateau. Acta Ecologica Sinica; 2004; 24(12): 2740–2756. https://doi.org/10.1088/1009-0630/6/5/011.

2. Daily GC. The value of nature and the nature of value. Science; 2000; 289: 395–396. https://doi.org/10.1126/science.289.5478.395.

3. Xie GD, Zhang CX, Zhang LM, Chen WH, Li SM. Improvement of the Evaluation Method for Ecosystem Service Value Based on Per Unit Area. Journal of Natural Resources; 2015; 30(8): 1243–1254. https://doi.org/10.11849/zrzyxb.2015.08.001.

4. Leemans HBJ, Groot RSD. Millennium ecosystem assessment: Ecosystems and human well-being: a frame work for assessment. PhysicsTeacher; 2003; 34(9): 534–534. https://doi.org/10.1119/1.2344558.

5. Costanza R, d’Arge R, de Groot R, Farber S, Grasso M, Hannon B, et al. The value of the world’s ecosystem services and natural capital. Nature; 1997; 387: 253–260. https://doi.org/10.1016/S0921-8009(98)00020-2.

6. Daily GC, Natures service: social dependence on natural ecosystems. Washington D C; 1997.

7. Sun J. Research advances and trends in ecosystem services and evaluation in China. Procedia Environmental Sciences; 2011; 10: 1791–1796. https://doi.org/10.1016/j.proenv.2011.09.280.

8. Zhang B, Li WH, Xie GD. Ecosystem services research in China: Progress and perspective. Ecological Economics; 2010; 69(7): 1389–1395. https://doi.org/10.1016/j.ecolecon.2010.03.009.

9. Shi Y, Wang RS, Huang JL, Yang WR. An analysis of the spatial and temporal changes in Chinese terrestrial ecosystem service functions. Chinese Science Bulletin; 2012; 57(17): 2120–2131. https://doi.org/CNKI:SUN:KXTB.0.2012-09-007.

10. Yu ZY, Bi H. The key problems and future direction of ecosystem services research. Energy Procedia; 2011; 5: 64–68. https://doi.org/5:64-68.10.1016/j.egypro.2011.03.012.

11. Ouyang ZY, Zheng H, Xiao Y, Polasky S. Improvements in ecosystem services from investments in natural capita. Science; 2016; 352(6292): 1455–1459. https://doi.org/10.1126/science.aaf2295.

12. Cao W, Li R, Chi X, Chen N, Chen J, Zhang H. Island urbanization and its ecological consequences: a case study in the Zhoushan Island,East China. Ecological Indicators; 2017; 76:1–14. https://doi.org/10.1016/j.ecolind.2017.01.001.

13. Li BJ, Chen DX, Wu SH, Zhou SL. Spatio-temporal assessment of urbanization impacts on ecosystem services: case study of Nanjing City, China. Ecological Indicators; 2016; 71: 416 –427. https://doi.org/10.1016/j.ecolind.2016.07.017.

14. Li L, Wang XY, Luo L, Ji XY, Zhzo Y, Zhao YC, et al. A systematic review on the methods of ecosystem services value assessment. Chinese Journal of Ecology; 2018; 37(4): 1233–1245. https://doi.org/10.13292/j.1000-4890.201804.031.

15. Boithias L, Terrado M, Corominas L, Ziv G, Kumar V, Marqués M, et al. Analysis of the uncertainty in the monetary valuation of ecosystem services-a case study at the river basin scale. Science of the Total Environment; 2016; 543(Pt A): 683–690. https://doi.org/10.1016/j.scitotenv.2015.11.066.

16. Richardson L, Loomis J, Kroeger T, Casey, F. The role of benefit transfer in ecosystem service valuation. Ecological Economics; 2015; 115: 51–58. https://doi.org/10.1016/j.ecolecon.2014.02.018.

17. Costanza R, Groot RD, Sutton P, Ploeg SVD, Anderson SJ, Kubiszewski I, et al. Changes in the global value of ecosystem services. Global Environmental Change; 2014; 26: 152 – 158. https://doi.org/10.1016/j.gloenvcha.2014.04.002

18. Xie GD, Lu CX, Zheng D, Li SC. Ecological assets valuation of the Tibetan Plateau. Journal of Natural Resources; 2003; 18(2): 189–196. https://doi.org/10.3321/j.issn:1000-3037.2003.02.010.

19. Xie GD, Zhen L, Lu CX, Xiao Y, Chen C. Expert knowledge based valuation method of ecosystem services in China. Journal of Nature Resources; 2008; 23(5): 911–919. https://doi.org/10.11849/zrzyxb.2008.05.019.

20. Boyle KJ, Bergstrom JC. Benefit transfer studies: myths, pragmatism, and idealism. Water Resources Research; 1992; 28(3): 657–663. https://doi.org/10.1029/91wr02591.

21. Loomis JB, Rosenberger RS. Reducing barriers in future benefit transfers: needed improvements in primary study design and reporting. Ecological Economics; 2006; 60(2): 343–350. https://doi.org/10.1016/j.ecolecon.2006.05.006.

22. Sharp R, Tallis HT, Rickett T. InVEST+VERSION+User’s guide. The natural capital project, Stanford university, university of Minnesota, the nature conservancy, and World Wildlife Fund [EB/OL]. (2013-03-05)[2018-04-05]. http://data.natura/capita/project.org/night/y-build/release_dufault/release_default/documentation/; 2013.

23. Sun X, Crittenden JC, Lif, Lu ZG, Dou XL. Urban expansion simulation and the spatio-temporal changes of ecosystem services,a case study, in Atlanta Metropolitan area, USA. Science of the Total Environment; 2018; 622: 974–987. https://doi.org/10.1016/j.scitotenv.2017.12.062.

24. Paracchini M L, Zulian G, Kopperoinen L, Maes J, Schaegner J P, Termansen M, et al. Mapping cultural ecosystem services: a framework to assess the potential for outdoor recreation across the EU. Ecological Indicators; 2014; 45(5): 371–385. https://doi.org/10.1016/j.ecolind.2014.04.018.

25. Ma FJ, Liu JT, Eneji AE. A review of ecosystem services and research perspectives. Acta Ecologica Sinica; 2013; 33(19): 5963 – 5972. https://doi.org/10.5846/stxb201306071398.

26. Chen Y, Li JF, Xu J. The impact of socio-economic factors on ecological service value in Hubei Province: A geographically weighted regression approach. China Land Science; 2015; 29(6):89–96. https://doi.org/10.13708/j.cnki.cn11-2640.2015.06.012.

27. Zhang C, Ren ZY, Gao MX, Yan WH. The forest ecosystem services and their valuation in Gansu Province. Journal of Arid Land Resources and Environment; 2007; 21(8): 147–151.

28. Wang SL, Liu XD, Wang JH, Li XB, Zhao ZQ. Soil conservation and value assessment of forest ecosystem in Gansu Province. Journal of Soil and Water Conservation; 2011; 5: 35–39. https://doi.org/CNKI:SUN:TRQS.0.2011-05-009.

29. Guo XN, Qi Y, Wang J, Liu BK, Wang FK, Chen ZH, et al. The research of forest ecosystem and their valuation of Gansu Province based on remote sensing and GIS. Remote Sensing Technology and Application; 2009; 2: 217–222. https://doi.org/10.11873/j.issn.1004-0323.2009.2.217.

30. Shu C, Zhang LB. Ecosystem service value assessment research based on GIS and remote sensing technology. Geomatics & Spatial Information Technology; 2015; 1: 30–32, 36. https://doi.org/10.3969/j.issn.1672-5867.2015.01.009.

31. Wang J, Qi Y, Chen ZH. Modeling dynamic assessment on ecosystem services based on remote sensing technology-A case study on Gansu grassland ecosystem. Journal of Glaciology and Geocryology; 2010; 6: 514–521. https://doi.org/10.3724/SP.J.1226.2010.00514.

32. Zhao HY, Chen Y, Yang J, Pei TT. Ecosystem service value of cultivated land and its spatial relationship with regional economic development in Gansu Province based on improved equivalent. Arid Land Geography; 2018; 4: 851–858. https://doi.org/10.12118/j.issn.1000-6060.2018.04.22.

33. De Groot RS, Alkemade R, Braat L, Hein L, Willemen L. Challenges in integrating the concept of ecosystem services and values in landscape planning, management and decision making. Ecological Complexity; 2010; 7(3): 260–272. https://doi.org/10.1016/j.ecocom.2009.10.006.

34. MA (Millennium ecosystem assessment). Millennium ecosystem assessment: Living beyond our means-natural assets and human well-being world resources institute, Washington DC; 2005.

35. Burkhard B, Krool F, Nedkov S, Müllera F. Mapping ecosystem service supply, demand and budgets. Ecological indicators; 2012; 21: 17–29. https://doi.org/10.1016/j.ecolind.2011.06.019.

36. Wang J, Wei YM, Sun XY. Effects of excessive grazing on grassland eco-system services valuation. Journal of Natural Resources; 2006; 21(1): 109–117. https://doi.org/10.3321/j.issn:1000-3037.2006.01.014.

37. Jiao L, Zhao CZ. The analysis and evaluation on grassland ecosystem service function value of Shandan Horse Field in Qilian Mountain National Nature Reserve. Journal of Arid Land Resources and Environment; 2013; 27(12): 47–52. https://doi.org/10.3969/j.issn.1003-7578.2013.12.009.

38. Li X, Ye YH, Wang YLT, Fu L, Yang MC, et al. Evaluation of grassland resource values in arid and semi-arid area-A case of Etuoke’s grassland resource balance sheet. Journal of Arid Land Resources and Environment; 2018; 32(5): 136–143. https://doi.org/10.13448/j.cnki.jalre.2018.153.

39. Li W, Zhao XS, Cui LJ, Sun BD, Lei YR, Ma QF, et al. Distribution and supply value of freshwater wetland resources in Jilin Province. Journal of Hydroecology; 2017; 38(1): 10–17. https://doi.org/10.15928/j.1674-3075.2017.01.002.

40. Xiao SC, Xiao HL, Lu Q, Li S, Wei H. Evaluation on China desert and sandy land ecosystem services based on Its related water process and regulating functions. Journal of Desert Research; 2013; 33(5): 1–9. https://doi.org/10.7522/j.issn.1000-694X.2013.00180.

41. Xue H. Study on ecosystem services of artificial system. Hangzhou: Zhejiang University; 2013.

42. Zhang XL, Xu ZJ, Zhang ZH, Wu LY. Environment purification service value of urban green space ecosystem in Qingdao City. Acta EcologicaSinica; 2011; 31(9): 2576–2584. https://doi.org/10.1111/j.1759-6831.2010.00113.x.

43. Gong W, Ou SH, Yang T, Yi CX. Quantitative evaluation of ecological value of urban green space based on GIS/RS. Journal of Anhui Agricultural Sciences; 2018; 46(3): 38–43. https://doi.org/10.13989/j.cnki.0517-6611.2018.03.016.

44. Duan YB, Lei YK, Wu BJ, Peng DD, Tian GX. Evaluation and dynamic study on the ecological service value for urban green space system in Zhengzhou. Ecological Science; 2016; 35(2), 81–88. https://doi.org/10.14108/j.cnki.1008-8873.2016.02.013.

45. Wang QH, Li JX, Peng Y, Lan WJ. Evaluation of service function of shrub ecosystem in China. Jiangsu Agricultural Sciences; 2019; 47(4): 233–237. https://doi.org/10.15889/j.issn.1002-1302.2019.04.053.

46. Cui XH. Valuation of terrestrial ecosystems services and values –A case study of desert ecosystems in China. Beijing: Chinese Academy of Forestry; 2009.

47. Zhao JH. Study on the influence of farmland irrigation mode on groundwater supply in Bayin River Basin. Xi’an: Chang’an University; 2018.

48. Xu T, Xu Y, Jiang B, Zhang L, Song WB, Zhou DM. Evaluation of the ecosystem services in caohai wetland, Guizhou Province. Acta Ecologica Sinica; 2015; 35 (13): 4295–4303. https://doi.org/10.5846/stxb201403250550.

49. Cui LJ, Zhao XS, Li W, Kang XM, Lei YR, Zhang MY, et al. Analysis on groundwater supply function of wetlands in Jilin Province based on soil permeability coefficient. Journal of Nature Resources; 2017; 32(9): 1457–1468. https://doi.org/10.11849/zrzyxb.20160905.

50. Guo LC, Yue P, Li HY, Xiang J, Ge P. Distribution and circular process of water resources in the arid area of Hexi Corridor. Agricultural Research in the arid areas; 2011; 29(6): 157–163. https://doi.org/CNKI:SUN:GHDQ.0.2011-06-032.

51. Zhang GH, Nie ZL, Wang JZ, Cheng XX. Isotopic characteristic and recharge effect of groundwater in the water circulation of Heihe river basin. Advances in Earth Science; 2005; 20(5), 511–518. https://doi.org/10.3321/j.issn:1001-8166.2005.05.005.

52. Jiang LZ, Yang DY, Mei CC, Ban YC, Yang XM. Ecosystem service function and its value assessment for urban green space-a case study on Futian District of Shenzhen City. Journal of HuaZhong Normal University (Natural Sciences); 2018; 3: 424–431. https://doi.org/10.19603/j.cnki.1000-1190.2018.03.020.

53. Cui CW, Xu XG. Ecosystem services assessment of the agricultural lands in Beijing region. Ecological Economy; 2007; 10: 338–340. https://doi.org/10.3969/j.issn.1671-4407.2007.10.093.

54. Xiao T, Gao YH, Zhai J, Yang M, Sun CX. Study on the value accounting of grassland ecosystem regulation service-a case study of Maduo County, Qinghai Province. Chinese Environmental Sciences Association; 2016; 650–655.

55. Mo FR, Chu XZ, Ma XF, Ma Q. The windbreak and sand fixation function and its value assessment of landscape patterns change of Ebinur lake wetland. Ecological Science; 2017; 36(6): 195–206. https://doi.org/10.14108/j.cnki.1008-8873.2017.06.027.

56. Zhao JL, Dong ZY, Chang ZF. Ecological compensation of a Desert water-Lifting irrigation project based on opportunity cost of ecosystem service value: A case study on Jingtaichuanwater-Lifting irrigation project. Arid Zone Research; 2019; 3: 743–751. https://doi.org/10.13866/j.azr.2019.03.26.

57. Wang YK, Guo SX, Wang J, Yuan H, Xu BL, Wang DR. Estimation of forest ecosystem service value in the Qilian Mountains National Nature Reserve in Gansu of China. Journal of Desert Research; 2013; 33(6): 1905–1911. https://doi.org/10.7522/j.issn.1000-694X.2013.00236.

58. Zhang HQ. Evaluation of forest ecosystem service value of Baishuijiang National Nature Reserve in Gansu Province. Lanzhou: Gansu Agricultural University; 2015.

59. Li C. Remote sensing estimation and spatiotemporal differentiation of land ecological service value in coastal saline alkali soil area. Shijiazhuang: Hebei Agricultural University; 2015.

60. Cheng HT. Evaluation of ecosystem service function of Baisha reservoir in Henan Province. Jilin agriculture; 2019; 14: 28. https://doi.org/10.14025/j.cnki.jlny.2019.14.006.

61. Chen HS. Evaluation of ecosystem service function of Lishimen reservoir in Zhejiang Province. Anhui Agricultural Science Bulletin; 2019; 25(1): 135–136. https://doi.org/10.3969/j.issn.1007-7731.2019.01.055.

62. Sun ZL, Li YN, Yu J, Wang FR. Ecosystem service evaluation of the six important reservoirs in Zhejiang Province. Journal of Zhejiang University (Science Edition); 2015; 42(3): 353–364. https://doi.org/10.3785/j.issn.1008-9497.2015.03.020.

63. Kong DS, Zhang H. Economic value of wetland ecosystem services in the Heihe National Nature Reserve of Zhangye. Acta Ecologica Sinica; 2015; 35(4): 972–983. https://doi.org/10.5846/stxb201305141053.

64. Jiang B, Chen YY, Rao EM, Zhang L, Ouyang ZY. Final ecosystem services valuation of Bosten Lake. Chinese Journal of Ecology; 2015; 34(4): 1113–1120.

65. Cheng Y, Liu PX, Bai Y, Ma YL, Pan JY. Analysis of temporal and spatial variation, driving factors and trend prediction of grain yield in Gansu Province. Agricultural Research in the Arid Areas; 2009; 27(4): 225–229.

66. Zhu ZG, Wang DY, Zhu LH, Hao YL. Correlation analysis farmland quality standard grain yield Binzhou City. Shandong Land Resources; 2015; 31(1): 75–79. https://doi.org/10.3969/j.issn.1672-6979.2015.01.019.

67. Wang YB, Hu YL, Bu CY, Mi, CL. Research on arable land grade haracteristics and spatial distribution in Gansu Province. Chinese Journal of Agricultural Resources and Regional Planning; 2017; 38(11): 138–144. https://doi.org/10.7621/cjarrp.1005-9121.20171119.

68. Zhang R, Yang H, Meng XJ, Wu HJ. Monitoring and evaluation on pratacultural status in Gansu. Grassland and Turf; 2017; 37(1): 1–7. https://doi.org/10.3969/j.issn.1009-5500.2017.01.001.

69. Tang HP, Tang SQ. Traits of water resources in Gansu Province and its conservation and utilization. Journal of Desert Research; 2000; 20(2): 213–216. https://doi.org/10.3321/j.issn:1000-694X.2000.02.021.

70. Padmavalli R Proc. Symposium resources development and planning. Geography Dept, Madras Univ; 1976; 1–19.

71. Zhang K. The influence of deforestation of tropical rainforest on local climate and disaster in Xishuangbanna region in China. Climatological Note; 1986; 35, 223–236. http://ir.xtbg.org.cn/handle/353005/4886.

72. Balling, Robert C. The impact of summer rainfall on the temperature gradient along the united states-mexico border. Journal of Applied Meteorology; 1989; 28(4): 304–308. https://doi.org/10.1175/1520-0450(1989)028<0304:TIOSRO>2.0.CO;2.

73. Meher-Homji VM. Probable impact of deforestation on hydrgical processes. ClimChande; 1991; 19: 163–173. https://doi.org/10.1007/bf00142223.

74. People’s Government of Gansu Province. The scope of groundwater over mining area, forbidden mining area and restricted mining area. http://www.gansu.gov.cn/art/2016/1/14/art_4785_261126.html; 2016.

75. Xie GD, Xiao Y, Zhen L, Lu CX. Study on ecosystem services value of food production in China. Chinese Journal of Eco-Agriculture; 2005; 13(3), 10–13. https://doi.org/CNKI:SUN:ZGTN.0.2005-03-003.

76. Duan RJ, Hao JM, Zhang JX. Land utilization and changes on eco-service value in different locations in Beijing. Transactions of the Chinese Society of Agricultural Engineering; 2006; 22(9): 21–28. https://doi.org/10.1111/j.1744-7917.2006.00098.x.

77. Wang SL, Liu XD, Wang JH, Li XB, Jin M, Zhang XL. Evaluation on forest ecosystem services value in Gansu province. Journal of Arid Land Resources and Environment; 2012; 3: 139–143. https://doi.org/10.1007/s11783-011-0280-z.

78. Qi ML. Land use change and influence to ecological service in Gansu Province. Journal of Anhui Agricultural Sciences; 2010; 38(20): 10814–10817. https://doi.org/10.3969/j.issn.0517-6611.2010.20.126.

